# Bacteriophage can promote the emergence of physiologically sub-optimal host phenotypes

**DOI:** 10.1101/621524

**Authors:** Hanna Schenk, Michael Sieber

**Affiliations:** Max Planck Institute for Evolutionary Biology, Plön, Germany

## Abstract

Reproduction of bacteria-specific viruses, or bacteriophage, requires the replication and translation machinery of the host cell. As a consequence, phage fitness depends intimately on the physiological state, i.e. growth rate, of the host. We include this dependence of critical phage traits on host growth rate in a mathematical model of a bacteria-phage interaction. This leads to a feedback loop between phage success, host population size, nutrient availability and host growth rate. We find that this feedback allows slow growing bacteria to have a competitive advantage in the presence of phage. Under certain conditions a slow growing host mutant can even drive the phage to extinction. Since in a phage-free environment slow growth is deleterious, the mutant subsequentely dies out as well, constituting a kind of altruistic scenario similar to abortive infections.

## Introduction

A parasite’s fitness is largely determined by the ability of a host to support parts or all of the infecting parasite’s life-cycle. This creates a strong dependence of the parasite’s success on host physiology (Bedhomme et al., 2004; Hall et al., 2009; Cornet et al., 2014). Crucially, host physiology can change dynamically in response to nutrient availability, which in turn may be influenced by parasite-induced changes in host population density. This creates the potential for an intricate feedback between host growth and parasite fitness, which is not commonly considered in theoretical models of host-parasite interactions. Therefore, linking parasite proliferation to the physiological status of the host and including physiologically diverse types increases the validity of models.

Viruses are good examples for obligate parasites which rely entirely on the host for reproduction. Bacteriophage, viruses that infect bacteria, are of particular interest since they act as model organisms in ecology and evolution and could serve as alternatives to antibiotics in medicine and agriculture (Altamirano and Barr, 2019). The life-cycle of lytic phage (Figure 1) can be broken down into a few crucial steps: First, phage adsorb to and inject their genetic material into the host cell. The host DNA is then degraded and the host’s protein synthesis machinery starts to produce the components which assemble into full phage particles. This process goes on until finally the hosts lyses and releases the new phage into the environment.

**Figure 1:**
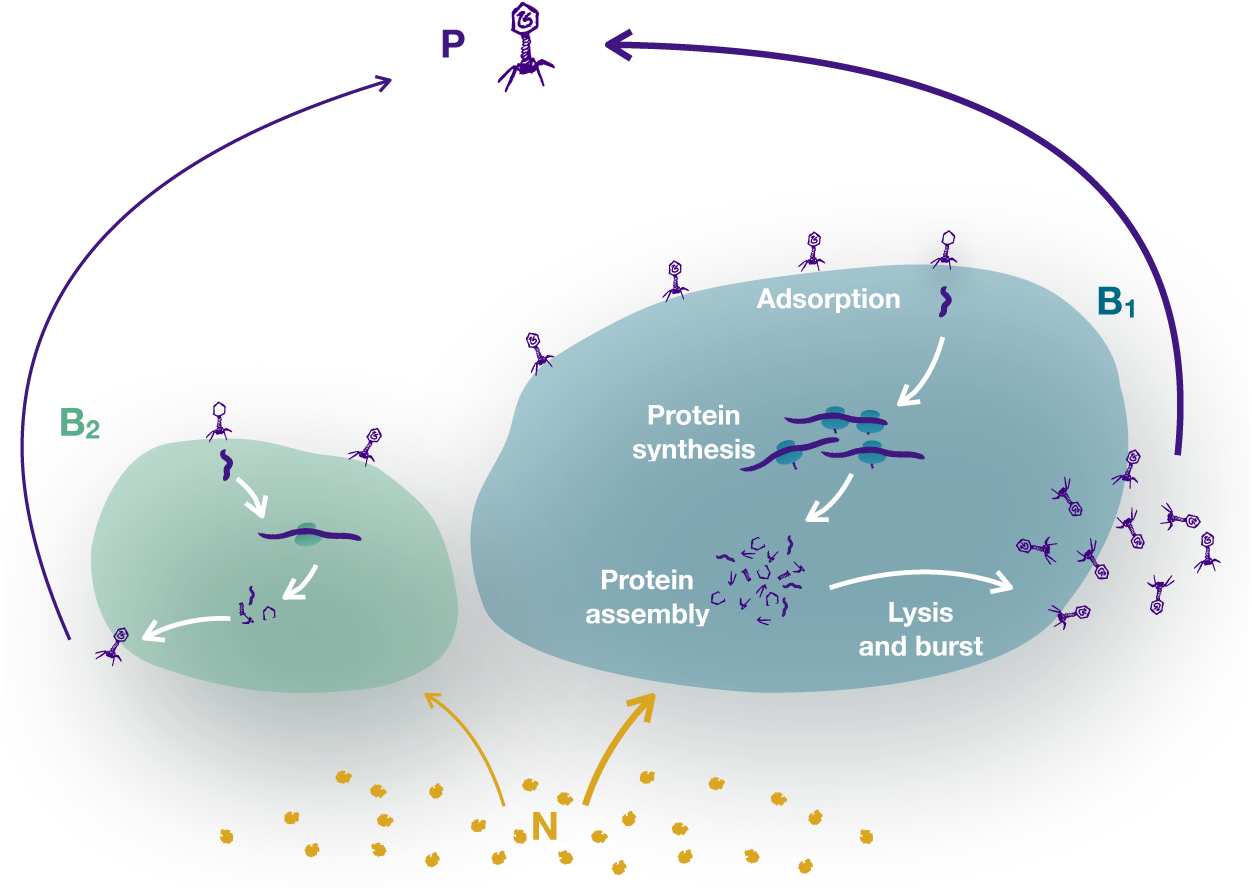
Lytic phage replication. depends on the physiology of the host: Slow growing bacteria are smaller. Fast growing bacteria are larger, making them an easier target for phage encounter. Fast growing cells have more machinery (ribosomes, resources), which can produce phage faster and leads to a larger burst size.

The rate of phage adsorption to host cells and the number of released phage after lysis, the burst size, have been shown to depend on the growth rate of the host (Hadas et al., 1997; Rabinovitch et al., 1999; You et al., 2002; Horas et al., 2018). In fact, below a certain growth rate of the host phage replication may cease completely (Rabinovitch et al., 2002; Golec et al., 2014). The addition or removal of specific nutritional compounds or the starvation of bacteria can influence the phage’s adsorption efficiency and duration and speed of intracellular phage replication (Propst-Ricciuti, 1976; Kokjohn and Sayler, 1991; Łoś et al., 2007; Briggiler Marcó et al., 2015; Zaburlin et al., 2017). A shorter cell doubling time increases the size of bacteria, therefore faster growing cells tend to have a larger cell surface area (Schaechter et al., 1958; Donachie, 1968; Vadia and Levin, 2015; Taheri-Araghi et al., 2015; Amir, 2017). There is great variation in the receptor usage of phage (Bertozzi Silva et al., 2016) and the details of adsorption depend on the number and spatial distribution of these receptors on the cell (Schwartz, 1976; Chatterjee and Rothenberg, 2012). But in general, a larger cell will provide a bigger target for phage (see differences in cell size in Figure 1), implying an increase of the adsorption rate with cell size (Delbrück, 1940; Hadas et al., 1997) up to a saturation point (Schwartz, 1976). After infection, the protein synthesising system of the host cell is allocated towards phage production. Therefore, a high number of ribosomes, found in faster growing and larger cells (see Figure 1), should in general speed up the production, thereby increasing the final burst size of phage (Hadas et al., 1997; Golec et al., 2014; Choua and Bonachela, 2019).

These dependencies can be formalized in two concrete statements about the entanglement of phage replication and host physiology: (i) The so-called bacterial growth law states that the cell volume, and thus surface area, increases exponentially with growth rate. Together with the linear increase of adsorption rate with cell surface area, the adsorption rate of phage depends exponentially on the growth rate of their hosts. (ii) Burst size is largely determined by the intracellular replication rate of the phage, which increases linearily with host growth rate, implying that the burst size increases approximately linearly with the host’s growth rate.

Burst size in particular can change by an order of magnitude depending on the state of the host population (Hadas et al., 1997; Rabinovitch et al., 1999). Despite this, these dependencies of phage parameters on the physiological state of the host are rarely incorporated into models of bacteria-phage interactions. Some notable exceptions are studies on phage adsorption kinetics (Moldovan et al., 2007; Smith and Trevino, 2009; Calsina and Rivaud, 2014) which improves the fit of theoretical models with experimental data (Santos et al., 2014). Further examples include explicit growth dependence in a mathematical model on viral evolution (Choua and Bonachela, 2019) and a reduced lysis capability at high bacterial densities (Weitz and Dushoff, 2008).

Here we investigate a mathematical chemostat model that relates adsorption rate and burst size to host growth rate. We allow for heterogeneity in the host population through phenotypes differing in their maximum growth rates. The resulting trade-off between growth and visibility to the phage allows for a coexistence of two bacterial types under phage infection. An in-silico experiment shows that slow growing bacteria can not only invade, but actually be advantageous to the resident host population by acting as a sink for phage particles. This can lead to the extinction of both the phage and slow growing bacteria, while the original bacterial host is rescued from phage infection.

## Mathematical model

We first introduce a bacteria-phage chemostat model with constant burst size *β* and adsorption rate *ϕ* based on the model presented in Levin et al. (1977). Nutrients with concentration *N*_0_ enter the chemostat at the constant rate *d* and remaining nutrients are washed out with the same rate *d*. Bacteria with density *B* take up and grow on those nutrients with growth rate *g* and conversion rate *c*. We choose a Monod growth rate 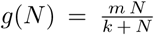 with half-saturation constant *k* which saturates at a maximum rate *m*. Phage with density *P* infect bacteria with the adsorption rate *ϕ*, which results in the loss of the infecting phage and the release of *β* new phage. The number *β* is called the *burst size*. The dynamics of nutrients, bacteria and phage over time (derivative 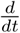 is denoted by a dot) are described by the following differential equations.

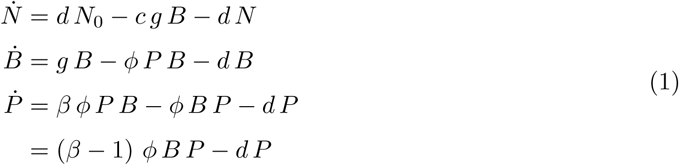

It is clear from the last equation that a necessary condition for phage survival is *β* > 1, reflecting that the burst size needs to at least offset the investment of the one phage required for the initiation of the infection process. The burst sizes of phage typically found in the literature range from a few dozens to thousands (De Paepe and Taddei, 2006), but these numbers are usually obtained when the hosts grow exponentially under optimal conditions. These conditions however, do not reflect the typical growth conditions of bacteria in the wild, where nutrients are often scarce and bacteria have to survive under starvation conditions for extended periods of time (Demoling et al., 2007; Aldén et al., 2001). Such conditions can be recreated in the chemostat with low nutrient inflow concentrations *N*_0_ and slow turnover rates *d*. While the model (1) takes into account the effect of changing nutrient levels on the physiology of the host via the Monod growth rate *g*(*N)*, this does not propagate to the burst size *β*, which is constant and does not change with the growth status of the host.

To explore the bacteria-phage dynamics when this simplifying assumption is relaxed, we now extend the standard model (1) by taking the dependency of the intra-cellular phage assembly rate on host physiology into account. More specifically, we incorporate the empirical observation that two key phage traits, the burst size *β* and the adsorption rate *ϕ*, depend on the current growth rate *g* of the host.

As phage replication relies completely on the host translation machinery, it is not surprising that the rate of this production depends on the host growth rate. Data from one-step growth experiments (Rabinovitch et al., 2002; You et al., 2002) suggests that if the host growth rate is not too high, the intra-cellular phage assembly rate is approximately a linear function of the host’s growth rate. The burst size can then be written as

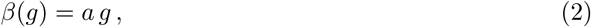

implying that it can now change dynamically, since the growth rate *g* = *g*(*N)* depends on the changing amount of nutrients *N* in the chemostat. Crucially, a low bacterial growth rate can now result in an effective burst size smaller than one, i.e.

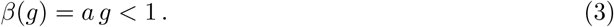

This corresponds to the paradoxical situation where infecting new host cells is actually detrimental to the phage, as the host is then effectively a sink for the phage.

The time it takes a single phage to find a host cell is determined by the density of the host in the environment and the adsorption rate *ϕ*, which will generally depend on the surface area of the host cell and the density of phage receptors on this surface. The density of receptors stays relatively constant at different growth rates (Hadas et al., 1997), while cell volume increases approximately exponentially with growth rate (Schaechter et al., 1958; Donachie, 1968; Taheri-Araghi et al., 2015; Amir, 2017). If for simplicity we assume cells as spheres with radius *r*, then the volume is *V* = 4*/*3*πr*^3^ *∝ e*^*g*^ and thus 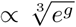. Since the density of phage receptors stays constant, the adsorption rate is proportional to the total cell surface area *ϕ ∝ A* = 4*πr*^2^ and thus we have 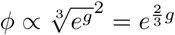. To keep the model as simple as possible, this function can be very well approximated by a quadratic function when *g* is not too large. Since adsorption rates saturate for high numbers of receptors (Schwartz, 1976), we further assume that this quadratic function also saturates at a maximum infection rate *q*, giving

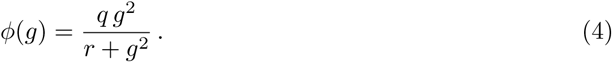

With these new key extensions, in our model a host’s maximum growth rate *m* not only defines its competitive ability in a phage-free environment, but also its value for an infecting phage. In this way host fitness and phage fitness become even more closely intertwined.

### Ecological long term behaviour

The system of ordinary differential equations (1) together with equations (2) and (4) has five fixed points where the rate of change in densities vanish 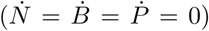. One is the trivial nutrients-only fixed point *N*\*= *N*_0_, which is unstable. A small amount of bacteria is enough to perturb the system to another fixed point with nutrient concentration 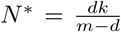 and bacterial density 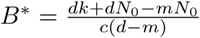. Three non-trivial coexistence points exist, two of which are unstable or negative for reasonable parameter combinations. Only one biologically relevant and locally stable coexistence point remains.

An important difference to models with a constant burst size is that phage can not easily invade a phage-free system and reach the coexistence equilibrium, since the bacteria-only fixed point is also locally stable (Fig. 2). This bistability has already been described by Weitz and Dushoff (2008), who in contrast to our more mechanistic model assume that lysis depends directly on bacterial densities via logistic growth. The reason for this bistability is that at the phage-free equilibrium nutrient concentrations are low and consequently the per-capita growth rate of the bacteria, which are at carrying capacity, is also very low. In this case the effective burst size is not large enough for the phage to grow to high densities, which would suppress the bacteria to lower densities where per-capita growth rates increase. If the initial phage population is large enough, however, lysis of cells will decrease bacterial densities enough to allow nutrients to reach higher concentrations, which in turn increases bacterial growth rate and phage burst size (Fig. 2).

**Figure 2:**
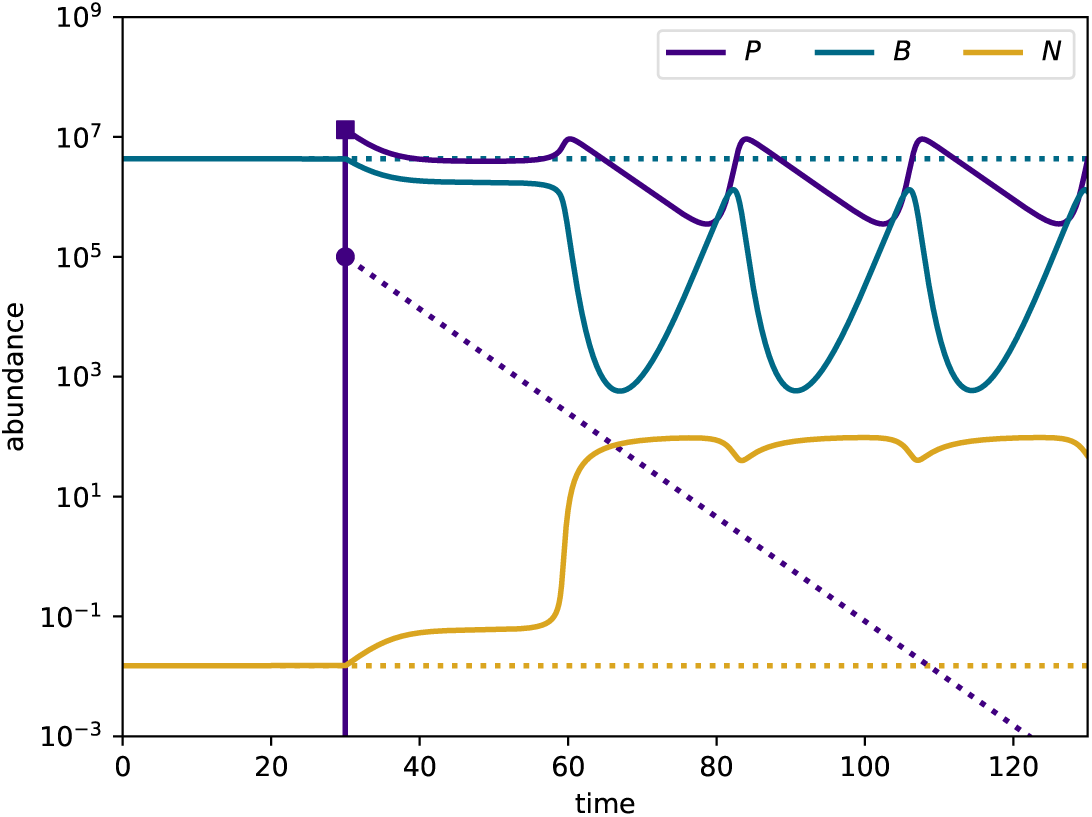
Example of bistability in model (1). Up to time point 30 the system is phage-free with bacteria at carrying capacity. The filled dot represents a small introduction of phage (10^5^ particles). Phage die out and nutrients and bacteria remain unchanged (dotted lines). The filled square and solid lines show an alternative scenario with a higher initial phage population (10^7^ particles). The bacterial population is suppressed and as a result the nutrient concentrations increase. This leads to a higher per-capita growth-rate of bacteria and higher burst sizes, allowing the phage to persist. Parameters: *q* = 5 × 10^−6^, *r* = 16, *k* = 0.07, *m* = 1, *a* = 1.2.

### Full model

We now extend system (1) by introducing two distinct types of bacteria, whose densities are denoted by *B*_1_ and *B*_2_. In the following we assume that those two types differ only in their maximal growth rates *m*_1_ and *m*_2_, and most of the time *B*_1_ will be the faster growing larger type (see Figure 1) with *m*_1_ ≥ *m*_2_. We then write 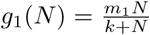 and 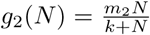 as shorthand for their respective growth rates, and similarly 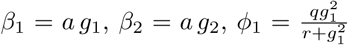 and 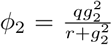 for the phage-related parameters. The implications of phenotypes with different maximal growth rates for the respective per-capita growth rates, burst sizes and infection rates is shown in Figure 3.

**Figure 3:**
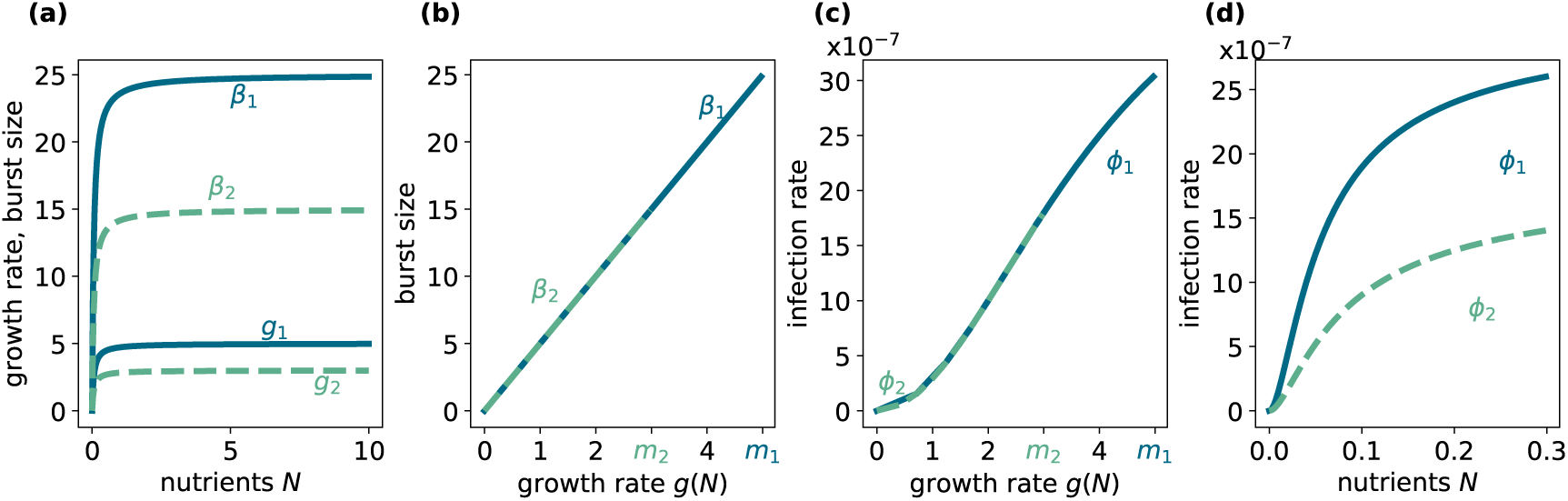
Growth rate, burst size and infection rate. Depending on the availability of nutrients *N* the growth rate *g* of bacteria can be at its maximum rate *m*_1_ (*m*_2_) or lower. Burst size *β* increases in the same way with nutrients (a), but linearly with the growth rate (b). Similarly, infection rate *ϕ*(*g*) increases nonlinearly with the growth rate (c) and therefore also with nutrients *N* (d). Parameters: *q* = 5 × 10^−6^, *r* = 16, *k* = 0.07, *m*_1_ = 5, *m*_2_ = 3, *a* = 1.2.

The full model with both bacterial types is then given by the differential equations

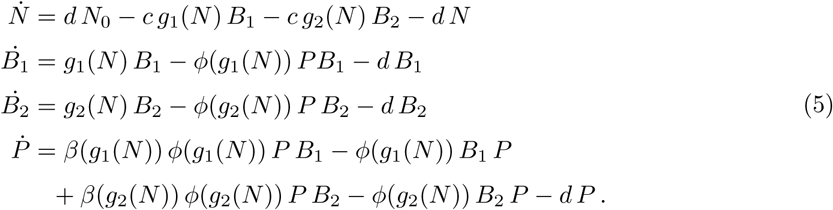

This system allows for additional fixed points compared to model (1). Either of two bacterial types can be in coexistence with the phage or inhabit the chemostat on its own (*N, B*). They cannot coexist without the phage when their growth rates are different, but with the phage coexistence becomes possible.It is challenging to determine the stability of the fixed points and thus the possible long term dynamics analytically, even when fixing some of the parameters. Therefore, in the following we employ an in-silico invasion experiment to study the dynamics, since this shows whether the fixed points are actually reached.

### In silico invasion experiment

In a phage-free environment, a slow growing bacterium will always lose against a fast growing but otherwise identical strain. When other selection pressures are at play other phenotypic properties might be more important and slow growth can become beneficial. We want to understand under which conditions an emerging slow growing mutant or migrant can take over the population of resident (fast growing) bacteria in the presence of phage.

To this end, we start an `in-silico` invasion experiment in a stable, phenotypically identical bacterial population of a resident type *B*_1_ which is infected by a phage *P*. This stable system is determined by the fixed point 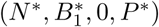 and we now consider what happens when a mutant *B*_2_ with a growth rate *g*_2_(*N)*, defined by *m* _2_, emerges in this system. The growth rate o f the resident *B*_1_ vanishes at the equilibrium and the mutant *B*_2_ needs a strictly positive per capita growth rate 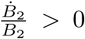 at the equilibrium to invade. The condition for initial mutant growth is therefore

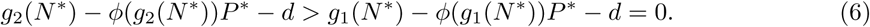

This invasibility condition is illustrated in Figure 4. Here, the value of the invader’s maximum growth rate *m*_2_ varies, while all other parameters are fixed, in particular the resident’s maximum growth rate *m*_1_ = 5. The figure shows the difference between the net per-capita growth rate of the invader (*g*_2_ – *d*) and the per capita death rate due to phage infections (*ϕ*_2_ *P*^*\**^). When the net growth is larger than the infection rate, the mutant can invade (shaded area). Note that there is a window of *m*_2_, in which the mutant can not invade. Above that window, the mutant is actually faster growing than the resident, and below, the infection rate decreases faster than the net growth.

**Figure 4:**
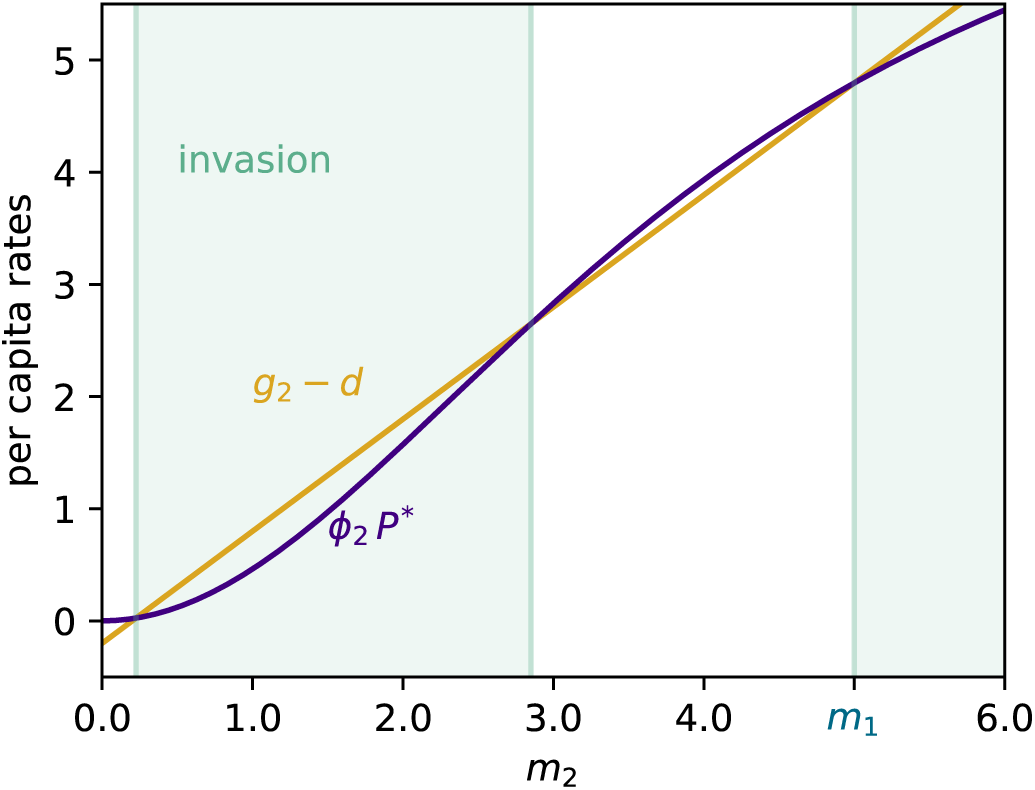
Initial invasion. Whether the mutant with maximum growth rate *m*_2_ can invade depends on the initial per capita growth rate, which is the net growth rate of the invader *g*_2_ – *d* minus death by phage infections *ϕ*_2_ *P*\*. Only if this difference, the overall per capita growth rate 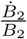, is positive can the slow growing mutant grow initially (shaded areas). See equation (6). Parameters: *m*_1_ = 5, *q* = 5 × 10^−6^, *r* = 16, *k* = 0.06, *c* = 2.3 × 10^−5^, *N*_0_ = 100, *a* = 5, *d* = 0.2.

In many cases, the small amount of invading slow growing bacteria can not overcome its disadvantage and immediately dies out again 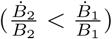. In the other cases, when Eq. (6) is fulfilled initially (shaded area in Figure 4), three scenarios are possible (cases A, B, C in Figure 5).

**Figure 5:**
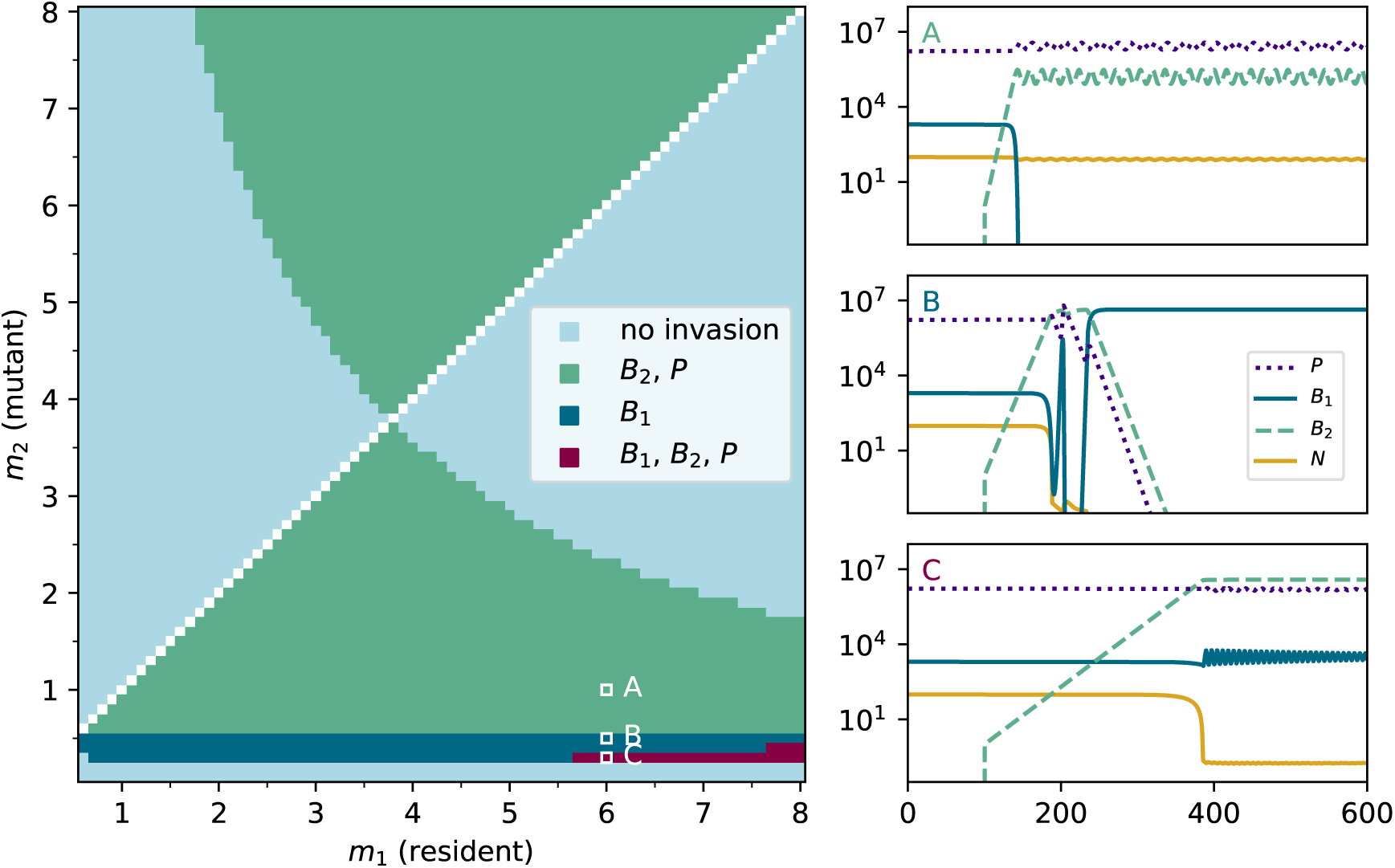
Invasion analysis. Pairwise invasion plot of a mutant in a stable resident-phage coexistence. Depending on how much slower or faster the mutant grows it can either die out immediately (light blue area) or invade the population. In the latter scenario three cases, detailed on the right are possible. Parameters: *q* = 5 × 10^−6^, *r* = 16, *k* = 0.06, *c* = 2.3 × 10^−5^, *N*_0_ = 100, *a* = 5, *d* = 0.2.

(**Case A**) The most common case is the replacement of the resident by the invader. In the course of the invasion the total bacterial load increases, which also increases phage abundance and slightly decreases nutrients. Whereas the slow growing and also smaller host *B*_2_ has a smaller surface area, the resident *B*_1_ is now infected with more phage and grows slightly slower. The effectively high infection rate drives the fast growing former resident to extinction. The coexistence of the slow growing *B*_2_ with phage is now possible, because generally *β*_2_ > 1. Here, phage can grow on the slower growing *B*_2_, albeit with a smaller burst size than on *B*_1_. Interestingly, due to a lower per-capita production of phage by the mutant *B*_2_, it reaches much higher densities than the previous resident, despite its lower growth rate. In this scenario phage densities remain essentially unchanged.

(**Case B**) A lower growth rate of the invader translates into a lower infection rate which results in high abundances of the invading *B*_2_. The total bacterial abundance is now so high that nutrients become extremely scarce. The resulting low growth rate of both hosts also reduces the burst size obtained from both hosts *β*_2_ < *β*_1_ ≈ 1. Phage numbers are reduced successively, which allows both hosts to become quite abundant temporarily. The result is the extinction of the phage and a transition to a completely phage-free system. Then, in the absence of phage, the faster growing *B*_1_ has a clear advantage and drives the slow growing bacteria *B*_2_ to extinction.

(**Case C**) A large difference between the growth rates of the fast growing resident and the slow growing invader, can lead to the rare coexistence case. The slow growing bacterium is very small and infected rarely (low *ϕ*_2_), and although it can grow to high abundances the growth rate remains small enough not to reduce nutrients as much as in case B. Thus, growth rates remain high enough to ensure burst sizes *β*_1_ > *β*_2_ ≈ 1 that sustain phage and a rare equilibrium, that maintains a relatively high abundance in both hosts, is reached.

In these particular examples we have observed the invasion of successively slower growing hosts. For an invasion analysis of both slow and fast invaders with maximum growth rates between 0.1 and 8, see the main pairwise invasion plot on the left in Figure 5. Here, we have chosen specific but reasonable parameters. Some parameters have been measured in (or applied to) chemostat experiments (*c, d*). In general, the scenarios also depend on all other parameters. For example the flow rate *d* within the chemostat also affects the invasion capability (equation (6)), the parameter *a* translates bacterial growth into phage burst and thus greatly changes the boundaries of the regions.

## Discussion

In this paper we incorporate the dependence of phage replication and adsorption on the host physiological state into a bacteria-phage chemostat model. In particular we consider two major relationships: (i) the number of phage progeny is linked to the host’s growth rate and (ii) the adsorption rate of the phage is determined by the size of the bacterial cell which increases with growth rate. In an *in silico* experiment, we probe the ability of a faster or slower growing bacterium to invade a population of resident bacteria in the presence of phage.

The scenarios encountered in the invasion experiment can be understood in terms of a trade-off between fast growth and two factors: (i) increased individual susceptibility to infection through larger cell size and (ii) increased population level spread of the infection because of higher burst sizes. Trade-offs are often included in theoretical studies (White and Bowers, 2005; Ehrlich et al., 2017), yet their mechanistic origins often remain obscure. In our model the trade-off arises naturally from the two assumptions. Fast growing bacteria are larger and therefore more visible to free floating phage resulting in an increased adsorption rate. The faster growing larger cells also have more replicative machinery, resulting in a larger phage burst size than slower and smaller cells. The shape of the trade-off and its magnitude change with the parameters of the system, but more importantly, it changes dynamically with the ecology of the system, since the nutrient-dependent growth rate changes in response to changes in host abundances, which in turn is affected by the intensity of phage infection.

A similar dependence of key phage traits on host physiology has recently been studied by Choua and Bonachela (2019), with a focus on the evolution of phage latent periods and without considering a feedback in physiological parameters of the host. Notably, Calsina and Rivaud (2014) made infection rate cell-size-dependent and additionally allowed adsorption on and ‘infection’ of already infected bacteria or dead cell material. This study introduced an age-structured model and proves co-existence of susceptible and resistant bacteria with phage when resistance comes with a trade-off in survival. The idea of including bacterial debris was inspired by Rabinovitch et al. (2003), who found that adsorption to debris stabilises a phage-bacteria coexistence in a simple recursion equation model.

It has been shown empirically (Bohannan and Lenski, 2000) and theoretically (Han and Smith, 2012), that when resistance is directly traded-off with growth, an invading resistant mutant does not cause the extinction of sensitive bacteria or phage. Previous studies have assumed a fixed gradual form of resistance, such as the gene-for-gene or modified gene-for-gene infection mechanisms (Forde et al., 2008). In contrast, here phage success on a particular host depends directly on its physiology (high or low maximum growth rate) and the dynamically changing ecological feedbacks in the host population. In this situation a third case beyond invasion or extinction becomes possible: invasion of a slow growing mutant, transition to a new stable phage-free state of the system, and subsequent extinction of the invader. When phage replication is slower than the rate of infection, a bacteria acts as an absorbing sink to the phage, like debris (Rabinovitch et al., 2003; Calsina and Rivaud, 2014). The sink provided by slow growing small bacteria can be counteracted by the production of phage in fast growing bacteria who now remain the only source of phage particles. Yet when nutrients are low, all bacteria become limited in growth and therefore limited in phage replication. If the depletion of phage by the sink is greater than the production by the source, or if there is no source due to limited growth the phage population will decline and go extinct, as in case B (Figure 5). After phage extinction, there is little benefit for the slow growing mutant on the level of the individual type, so it dies out and the system settles in a host-only equilibrium which is stable against invasion by either slow growing bacteria or phage. This is a kind of ecological hysteresis, where a change in the host population (emergence of slow growing phenotype) sends the system over a tipping point (extinction of the phage), from which the system does not return to its initial state, even after the inducing agent has gone extinct as well. This behaviour is caused by a general effect that phage have on their host populations, which only comes to light when the interdependence of host physiology and phage replication is taken into account. Generally, the phage suppresses its host population well below its carrying capacity, resulting in abundant resources and continued high per-capita growth and fast host turnover. As soon as phage abundance declines, however, the host recovers and as it approaches carrying capacity, its per-capita growth rate falls below the minimum required for a sustainable effective burst size. This prevents the re-infection by phage, but crucially only if the dependence of the burst size on host growth rate is taken into account. Thus, initial exponential growth in the mutant (Figure 4) does not imply a takeover of the population. The related bistability (Figure 2) of the phage-bacteria coexistence point has major implications for phage therapy, since it shows that the amount of phage delivered to the site of a bacterial infection is crucial for determining the success of the treatment.

By modulating the effective burst size of the phage, host cell physiology can be considered a plastic post-adsorption resistance mechanism on the population level. Such post-adsorption resistance mechanisms can limit phage host ranges (Sieber and Gudelj, 2014) and our results here show that physiological changes, such as lower growth rates, can lead to a very similar effect. More generally, this modulation of the effective burst size through changes in host growth rate shares similarities with abortive infections, where the host cell blocks phage multiplication and triggers cell death (Labrie et al., 2010). This altruistic behaviour of the infected cell limits phage replication and spread to other cells, much like the growth-rate mediated reduction in burst size we have considered in our study. Although we have chosen the specific example of bacteria and phage, parasites always depend to some extent on the physiological status of their host. As we have shown this opens up additional routes for the host to counteract parasite infections.

